# Contextual expectations shape cortical reinstatement of sensory representations

**DOI:** 10.1101/2021.08.05.455275

**Authors:** Alex Clarke, Jordan Crivelli-Decker, Charan Ranganath

**Affiliations:** Department of Psychology, University of Cambridge, UK; Center for Neuroscience, University of California, Davis, USA; Department of Psychology, University of California, Davis, USA

## Abstract

When making a turn at a familiar intersection, we know what items and landmarks will come into view. These perceptual expectations, or predictions, come from our knowledge of the context, however it’s unclear how memory and perceptual systems interact to support the prediction and reactivation of sensory details in cortex. To address this, human participants learned the spatial layout of animals positioned in a cross maze. During fMRI, participants navigated between animals to reach a target, and in the process saw a predictable sequence of five animal images. Critically, to isolate activity patterns related to item predictions, rather than bottom-up inputs, one quarter of trials ended early, with a blank screen presented instead. Using multivariate pattern similarity analysis, we reveal that activity patterns in early visual cortex, posterior medial regions, and the posterior hippocampus showed greater similarity when seeing the same item compared to different items. Further, item effects in posterior hippocampus were specific to the sequence context. Critically, activity patterns associated with seeing an item in visual cortex and posterior medial cortex, were also related to activity patterns when an item was expected, but omitted, suggesting sequence predictions were reinstated in these regions. Finally, multivariate connectivity showed that patterns in the posterior hippocampus at one position in the sequence were related to patterns in early visual cortex and posterior medial cortex at a later position. Together, our results support the idea that hippocampal representations facilitate sensory processing by modulating visual cortical activity in anticipation of expected items.

## Introduction

Our knowledge of how the world is structured has a powerful influence on our perceptions (Bar, 2004; de Lange et al., 2018; Oliva and Torralba, 2007; Summerfield and Egner, 2009). This is illustrated across a diverse range of phenomena, including visual illusions such as the Kanizsa triangle and enhanced object recognition when items appear in the correct spatial context (Davenport and Potter, 2004; Munneke et al., 2013; Palmer, 1975; Võ et al., 2019), which indicates that information we learn through experience can alter perceptual processes, and allow predictions about future states we may encounter (Bar, 2009; Eichenbaum and Fortin, 2009). Perceptual expectations, or predictions, can take different forms, with one useful distinction being between predictions of future states dependant on temporal continuity (such as a train moving from one location in space to another as it’s driven), and predictions of future states relating to upcoming items or events that are not currently in view (such as when turning a corner while navigating a familiar route), thus relying on memory processes and learned contextual associations. This latter form of perceptual expectation must depend on coordinated responses between perceptual and memory systems, however the neural implementation of prediction in this regard is elusive, especially in terms of how memory systems might moderate expectations to reactivate relevant perceptual details in cortex.

Expecting to see a specific visual stimulus generates responses in early visual cortex that resembles activity when perceiving the same stimulus, with suggestions that expectations result in stimulus templates being evoked in primary sensory regions (Kok et al., 2014, 2012). This premise is additionally supported by work showing that when items are expected, but not actually shown, activity in visual areas is similar to if the item did appear, despite the absence of any visual input (Eagleman and Dragoi, 2012). Such evidence is taken to suggest that stimulus representations in visual areas are sharpened by expectations, through a process of supressing activity that is inconsistent with the expectation (de Lange et al., 2018).

To make accurate predictions requires that we have previously encountered similar situations and learned what is likely to occur next. Statistical learning approaches argue that we acquire knowledge about how the world is structured through repeated experiences (Sherman et al., 2020; Shapiro et al., 2017), with this learned information utilised for navigation and guiding predictions of future states (Barron et al., 2020; Stachenfeld et al., 2017; Turk-Browne, 2019). In humans, the hippocampus has been shown to represent predictions of future states (Brown et al., 2016; Eichenbaum and Fortin, 2009), in addition to representing learned sequences of visual events in a distinct manner (Ezzyat and Davachi, 2014; Hsieh et al., 2014), together suggesting the hippocampus might be a primary source of top-down predictions that could result in the reactivation of expected sensory details in cortex. This is supported by recent evidence from Kok and Turk-Browne (2018), who showed that when an auditory cue is predictive of a subsequent visual item, hippocampal responses reflected the predicted stimulus and visual regions reflected the perceived stimulus. Such studies are beginning to reveal how memory and perceptual systems reflect aspects of prediction and perception, suggesting that the hippocampus has a top-down effect on future states in cortex. It is clear that the learned prior context must guide the prediction of future states, however direct empirical support for this contextually dependent hippocampal-cortical interaction remains limited.

While much of the focus has been on primary sensory regions and the hippocampus, a network of posterior brain regions have also been implicated in processing item and context-based reactivations and expectations, including the parahippocampal cortex, precuneus/posterior cingulate cortex and angular gyrus (Bar and Aminoff, 2003; Caplette et al., 2020; Jonker et al., 2018; Lee and Kuhl, 2016; Livne and Bar, 2016; Long and Kuhl, 2021). For instance, Capalette et al., (2020) compared activation when an object could be predicted following a visual scene, against trials when the object could not be predicted, finding that the precuneus was more active when predictions could be made, suggesting that the region may play a role in integrating contextual and object details. These posterior regions comprise a posterior medial (PM) network, implicated in processing contextual and episodic details (Ranganath and Ritchey, 2012; Ritchey and Cooper, 2020), which displays connectivity with the posterior hippocampus (Barnett et al., 2021, 2019). In contrast, the anterior hippocampus has connectivity with an anterior temporal (AT) network including the perirhinal cortex, temporal pole, amygdala and orbitofrontal cortex, and is thought to represent item and object information, but not contextual details (Ranganath and Ritchey, 2012; Ritchey and Cooper, 2020). This differing connectivity profile of the hippocampus has been linked to differential functional properties of anterior and posterior hippocampus (Poppenk et al., 2013), leading us to predict that context and contextually-driven reactivations will be present in the posterior hippocampus and PM network, in addition to the reactivation of sensory details in primary visual cortex. A further unexplored question is how the hippocampus, PM network and visual cortex interact to support the prediction and reactivation of sensory patterns in cortex.

To explore these issues, we conducted an fMRI study where participants navigated through a learned space and saw predictable sequences of objects. The learned space was comprised of objects positioned in a cross-maze, which was used to create sequence trials containing the items that would be seen when navigating from one end-point to another (Figure 1AB). Critically, a quarter of the sequence trials terminated early, where a participant expected to see a specific item given the past items, but instead a blank screen was presented. These catch trials resulted in activity patterns driven only by the expectations without any bottom-up visual input. Using multivariate pattern similarity analysis, our design allows us to ask (1) which regions represent information about the currently viewed item in the sequence context, (2) are these item representations specific to a given sequence context, and (3) do regions reflecting item information also represent information about the expected item, revealed using the catch trials. To more specifically test how the hippocampus interacts with other regions to enable such predictions, we employed multivariate representational connectivity (Anzellotti and Coutanche, 2018; Kriegeskorte et al., 2008; Pillet et al., 2020) to test whether the representational similarity structure in the hippocampus at one point in the sequence related to the structure in cortical regions at a later point in the sequence, which would provide evidence that hippocampal representations relate to the reactivation of expected future activity patterns in cortex.

**Figure 1.**
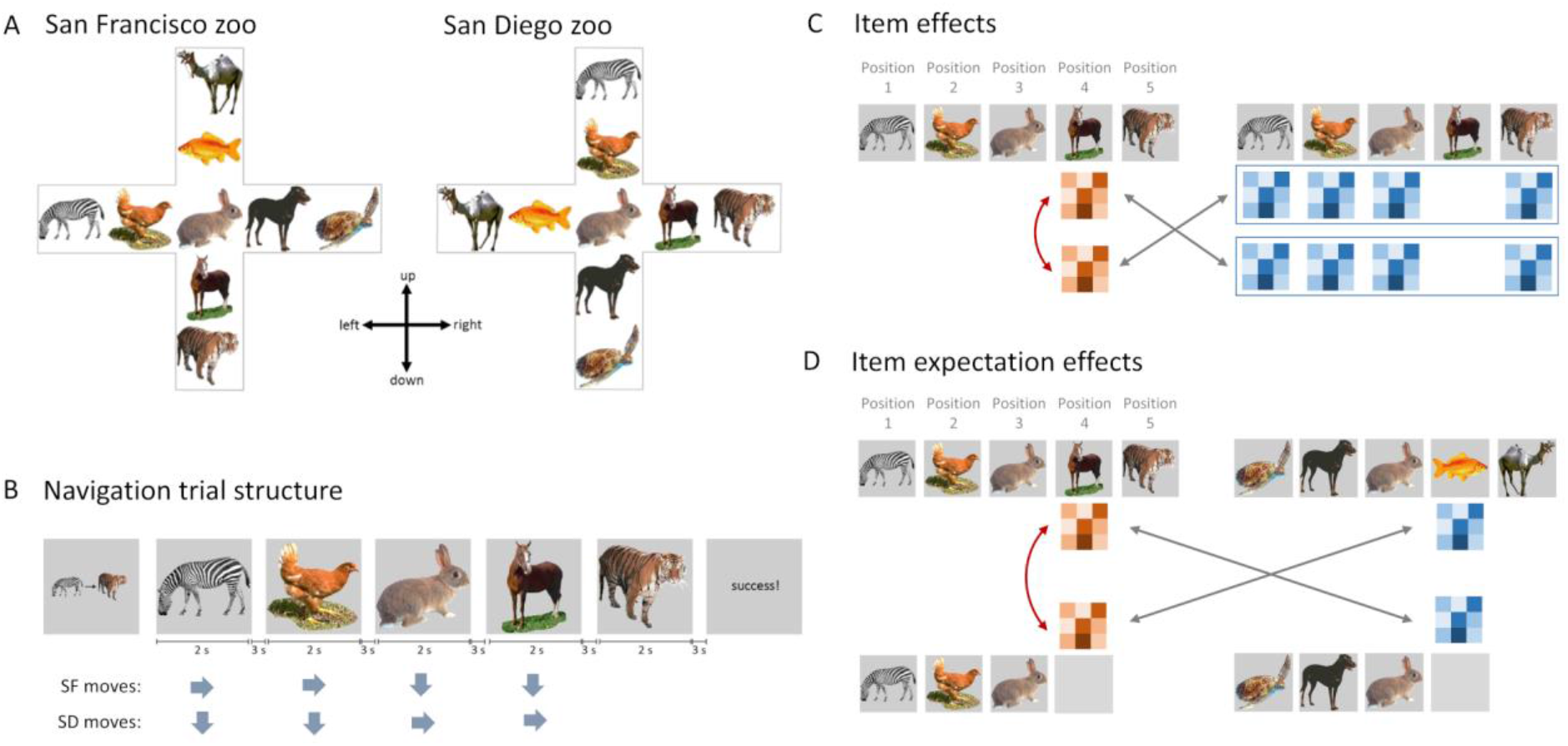
Experimental design and analysis. A) Participants learned the spatial layout of 9 animals in two distinct zoos. B) During fMRI, participants navigated between two animals, seeing a predictable sequence of images. C) Item effects were established by comparing activity patterns when seeing the same item in the same sequence and same zoo (red arrow), with patterns for different items within the same sequence context and zoo (grey arrows). D) Item expectation effects were established by comparing activity patterns between the blank period (catch trial) and the item that was expected given the sequence context (red arrow), against activity patterns for different blank and seen items (grey arrows).

## Results

### Behavioural learning and task performance

The experiment was conducted in two parts, a pre-fMRI learning session and a sequence navigation task during fMRI. During the pre-fMRI session, participants were required to learn the identities and locations of nine different animals in two related zoos (Figure 1A). The same animals were found in both zoos and animal transitions were maintained by mirror-reversing and rotating 90 degrees anti-clockwise one map to create the other. This ensures that the same sequences of animals could be seen in both zoos. During the pre-fMRI learning session, participants completed various behavioural tasks to learn the animals and their locations within each zoo. Participants completed map creation, exploration and navigation trials for each zoo, before a final behavioural test requiring navigation trials for both contexts. Navigation trials required participants to make a series of five moves between a start and goal animal, where start and goal animals were always end points of a maze arm (Figure 1B). Only the current-position animal was shown, and making the correct response resulted in seeing the next animal image in sequence. Correctly navigated trials therefore resulted in seeing a sequence of five animals in a predictable order. Participants were required to meet criterion performance (85% correct) before beginning the scanning session on the same day.

During the fMRI scanning session, participants completed six runs of navigation trials, where each run consisted of blocks of eight trials from each zoo. In each zoo, there were 12 different navigation trials, with each full sequence of five animal items being repeated three times. As the same animals were found in both zoos, with the same transitions, identical visual sequences were seen in both zoos. During scanning, participants showed a high level of performance for both zoos (San Francisco: mean = 94.3%, SD = 6%, San Diego: mean = 95.0%, SD = 5%) with no statistical differences seen between them (t(22) = 1.11, p = 0.28). This high level of performance during both learning and scanning indicates that the sequences were well-known and therefore participants would be able to predict what animal was to appear next (although they were not instructed to do this).

### Item, sequence and expectation effects in the early visual cortex and hippocampus

Our initial analysis of the fMRI data focussed on the early visual cortex and the hippocampus, both of which are suggested to support sensory expectations and predictions (e.g. Hindy et al., 2016; Kok and Turk-Browne, 2018). Given the different connectivity profiles, we additionally divided the hippocampus into a posterior and anterior section (Poppenk et al., 2013). Using multivariate pattern similarity analysis, we tested the extent to which these regions represented information about: (1) the currently viewed item, (2) the specific sequence context, and, (3) the next item that was expected in the sequence.

On each scanning navigation trial, participants made a series of five moves between a start and a goal animal, seeing a sequence of five animals. To address our questions at the level of individual items, fMRI single-trial beta values were estimated for each item in each sequence. Our analyses focussed on the item in position 4 of the sequence, first because position 4 items are always preceded by the same image in all sequences, meaning any impact of the preceding item on voxel patterns due to autocorrelation is controlled for. Second, one quarter of position 4 trials were omitted with the trial ending early, allowing us to study the impact of expectations through these catch trials. Third, position 4 is also situated after a key decision point (position 3), where it is possible to see multiple different animals following the central item (the rabbit), with the decision made at this point determines the next image. Therefore, position 4 allows us to both examine item-level effects, while controlling for autocorrelation and recent visual effects, and predictive effects generated from the central decision point.

We first determined if early visual cortex and the hippocampus were sensitive to information about the currently viewed item. To do this, we contrasted voxel pattern similarity (PS) for repetitions of the same item and compared this to PS between different items (Figure 1C). In order to control for sequence effects, PS was restricted to trials from the same sequence context and zoo, meaning that we are asking whether the items are dissociable within a specific sequence. Significant item effects were seen in V1/V2 (mean = 0.052, t(22) = 5.26, p < 0.0001) and the posterior hippocampus (mean = 0.011, t(22) = 2.02, p = 0.028), but not the anterior hippocampus (mean = 0.010, t(22) = 1.37, p = 0.09; Figure 2A).

**Figure 2.**
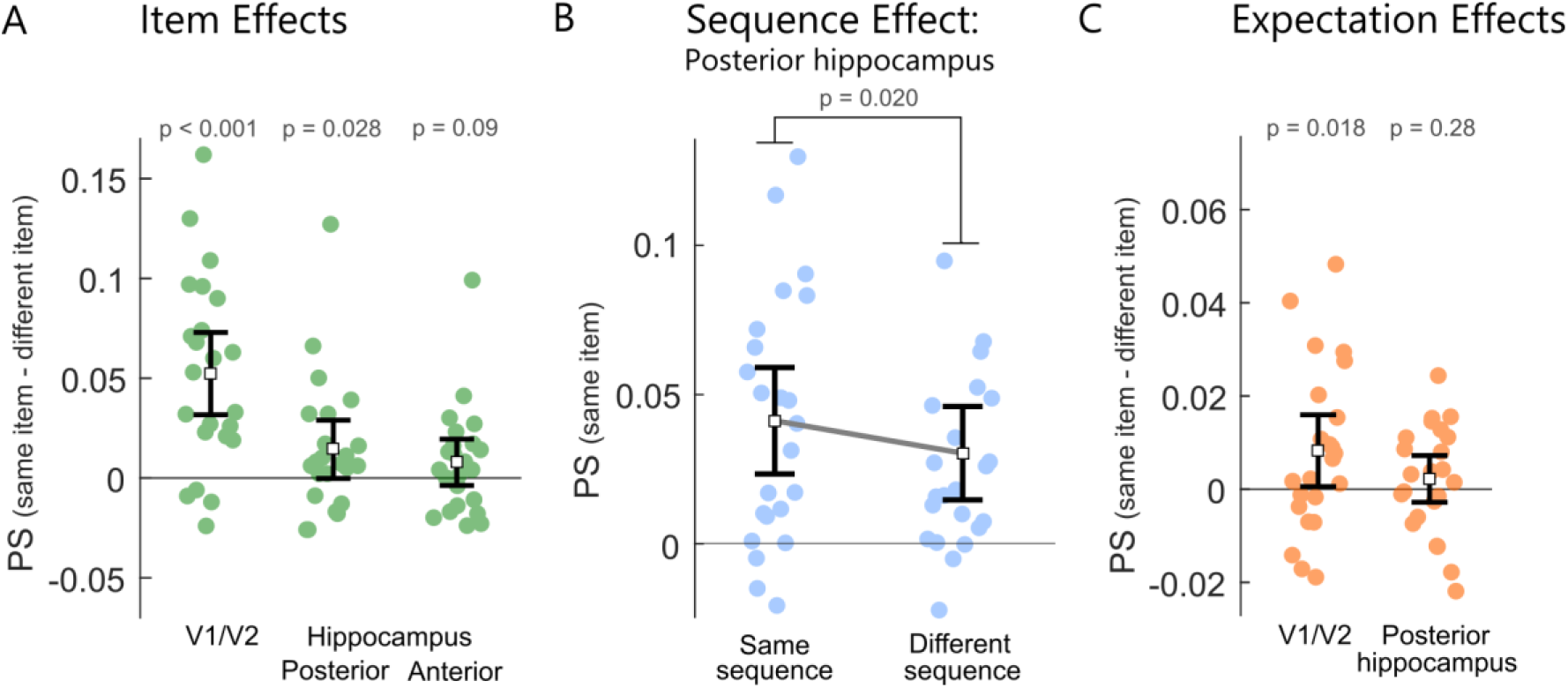
Pattern similarity results for early visual cortex and the hippocampus. A. Item effects showing difference in pattern similarity between same item pairs and different item pairs. B. Sequence effects in the posterior hippocampus showing changes in pattern similarity for same item pairs according to the sequence. C. Item expectation effects showing the difference in pattern similarity when the same item was omitted or seen, and when the omitted and seen item were different. Error bars show 95% confidence intervals around the mean.

These item-level representations may reflect the visual appearance of the object, however, given past research showing the hippocampus is sensitive to contextual and sequence information (Ezzyat and Davachi, 2014; Hsieh et al., 2014), we next asked if these item effects were dissociable across the different sequences. Here, we are interested in the sequence-level contexts, and not potential global zoo differences, meaning our analyses focussed on differences between the sequences without considering the two zoos. We compared PS between the same-items in the same sequence, to PS for the same-items when found in different sequences. As our analysis only included items in position 4, and PS is calculated between same items pairs, any differences we see are driven purely by information pertaining to the sequence, and not by visual details of item or temporal order. Significant sequence effects were found in the posterior hippocampus, where patterns were more similar for same-items from the same sequence (mean = 0.041) compared to same-items across different sequences (mean = 0.030; t(22) = 2.52, p = 0.0196). No significant sequence effects were observed in V1/V2 (t(22) = 1.81, p = 0.085) or the anterior hippocampus (t(22) = 1.22, p = 0.23). These results show that the posterior hippocampus not only represents information about the current item, but that these representations are further reflective of the specific sequence the item occurred in.

In the above analysis, we characterized representations of presented objects within a learned sequence. In well-learned sequences, upcoming items are known, and according to predictive models of perception, being able to predict upcoming items should impact neural processing by generating expectations about what is about to happen (Bar, 2004; de Lange et al., 2018; Trapp and Bar, 2015; Turk-Browne, 2019). To test this hypothesis, we focused our next analyses on catch trials (see Figure 1D), in which each sequence was terminated early, such that, after position 3, a blank screen was shown for six seconds, followed by the onset of the next navigation trial. In other words, on every catch trial, there was no motor response or external visual stimulation, and the preceding image was matched for all sequences. Thus, any representational content during a catch trial would be expected to be driven by memory-driven predictions in the absence of bottom-up input.

To test whether the regions that were sensitive to perceived items also carried information about expected items in the absence of sensory input, we assessed PS between presented items and activity when an item was expected but omitted from the sequence. PS was calculated between items and the catch trials when they were the same item, and compared to when the presented and catch trials were different (Figure 1D). Significant item expectation effects were seen in V1/V2 (mean = 0.008, t(22) = 2.25, p = 0.018) but not the posterior hippocampus (mean = 0.002, t(22) = 0.58, p = 0.28; Figure 2C). Our analyses clearly show that activity patterns in early visual regions are not only shaped by the bottom-up visual input, but that contextually-predicted item information is reactivated which matches the expected visual input.

Together, our analysis of early visual cortex and the hippocampus reveals that while item information is present in both regions, only item representations in the posterior hippocampus were modulated by the sequence the item was in, and only in the early visual cortex did we find evidence of item patterns being reactivated when they were expected, but failed to appear.

### Item and expectation effects in the posterior medial cortex

As discussed earlier, a wider network of regions beyond the hippocampus have been implicated in memory-guided predictions and contextual reactivations (Bar and Aminoff, 2003; Caplette et al., 2020; Jonker et al., 2018; Lee and Kuhl, 2016; Livne and Bar, 2016; Long and Kuhl, 2021). The PM network is functionally connected to the posterior hippocampus, and associated with reactivation of contextually-relevant object information, whilst the AT network is connected to the anterior hippocampus and is thought to represent item information (Ranganath and Ritchey, 2012). As such, we next repeated our analysis of item, sequence and expectation effects across regions in the PM network - parahippocampal cortex (PHC), posterior medial cortex (PMC; precuneus/posterior cingulate cortex) and the angular gyrus - and the AT network - the temporal pole and perirhinal cortex (PRC).

Item effects were calculated by comparing PS between same-item pairs with PS for different item pairs, within the same sequence. Significant item effects were seen in PMC (mean = 0.026, t(22) = 3.11, p = 0.0128) and the PHC (mean = 0.016, t(22) = 2.80, p = 0.013), but not the angular gyrus (mean = 0.010, t(22) = 1.44, p = 0.10), temporal pole (mean = 0.11, t(22) = 1.74, p = 0.080) or PRC (mean = 0.005, t(22) = .95, p = 0.17; Figure 3A). This suggests that in addition to the early visual cortex and posterior hippocampus, regions of the PM network – the PMC and PHC - also represent the currently viewed item during navigation.

**Figure 3.**
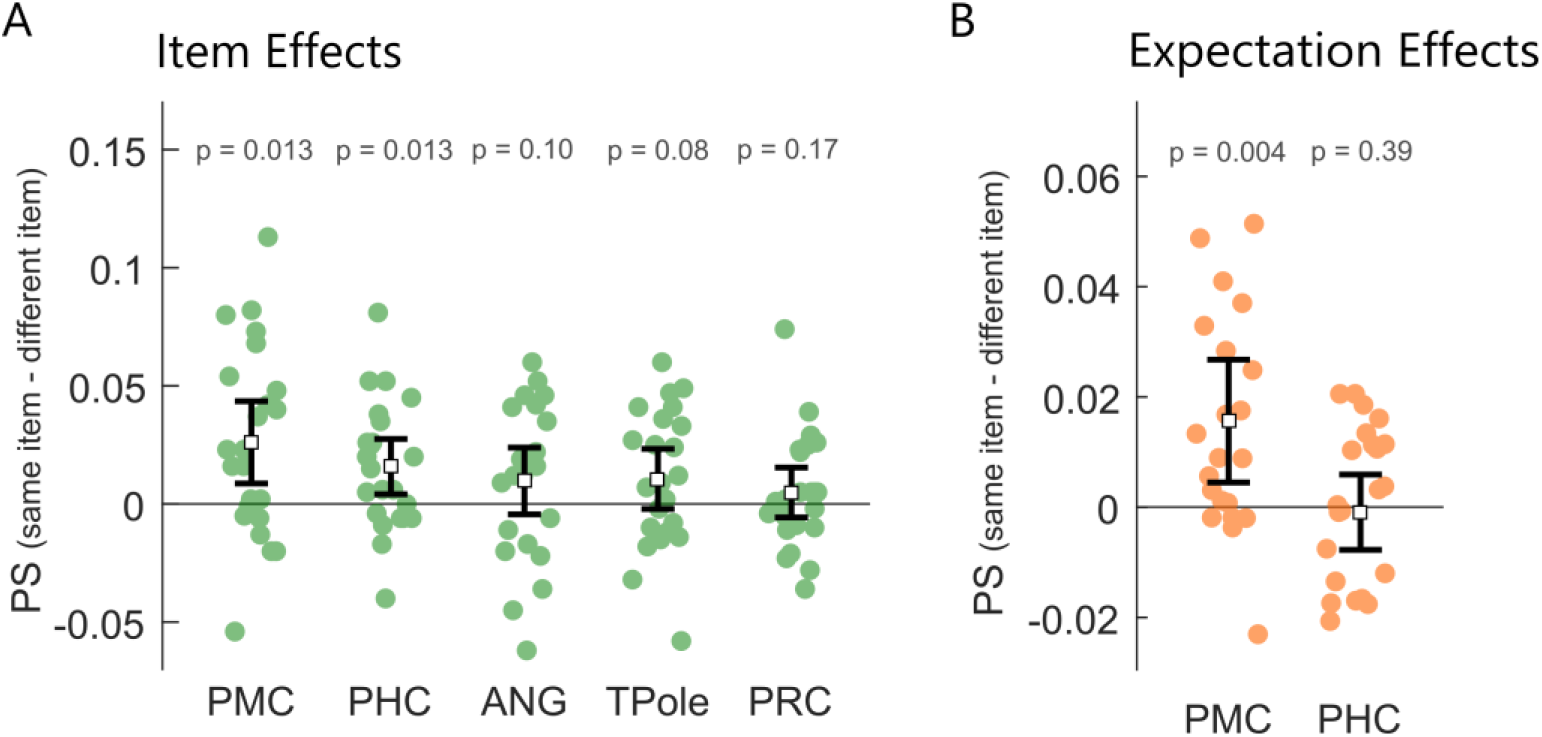
Pattern similarity results for regions of the PM and AT networks. A. Item effects showing difference in pattern similarity between same item pairs and different item pairs. FDR correction applied to p-values. B. Item expectation effects showing the difference in pattern similarity when the same item was omitted or seen, and when the omitted and seen item were different. Error bars show 95% confidence intervals around the mean.

We next asked whether these regions represented the same items in a distinct manner across sequences by comparing PS for same-item pairs from the same sequence, against same-item pairs across different sequences. This analysis revealed no significant sequence effects (all p’s > 0.05, fdr corrected). Finally, we tested whether the regions that showed item effects also showed effects of expected items in the absence of bottom-up visual input by comparing PS for when items and catch trials were the same item, to when the presented and catch trials were different. Significant item expectation effects were in the PMC (mean = 0.016, t(22) = 2.96, p = 0.0041) but not the PHC (mean = −0.001, t(22) = 0.27, p = 0.39; Figure 3B).

In addition to hypothesis-driven ROI analyses, we further conducted an exploratory searchlight analysis to identify other potential areas carrying information about perceived and predicted stimuli. Results from this analysis, reported in the Supplementary Information, showed strong item and expectation effects in posterior aspects of the visual cortex, consistent with our findings in the V1/V2 ROI analyses.

Overall, our data point to a representation of the current item in a network of regions in early visual, PM and the posterior hippocampus, with item representations in the posterior hippocampus being further specific to the sequence context. Crucially, representations in early visual cortex and the PMC reflect the expected item in the absence of any bottom-up input, suggesting they are a site of top-down effects based on contextual expectations.

### Hippocampal and cortical interactions support reactivation

An important question arising from our results, is by what mechanism are visual details of the items being reactivated in early visual cortex and PMC? Motivated by the role of the hippocampus in representing sequence knowledge, cortical reinstatement and the prediction of future states (Eichenbaum and Fortin, 2009; Hindy et al., 2016; Kok et al., 2019; Stachenfeld et al., 2017; Turk-Browne, 2019), we next tested the hypothesis that hippocampal pattern information related to future information states in visual cortex and PMC - regions showing item expectation effects, which may suggest the hippocampus supports the prediction and reactivation of sensory patterns in cortex. We reasoned that, if this is the case, the fidelity of a hippocampal sequence representations at one state (as indexed by PS across same-sequence pairs) should be predictive of the fidelity of the cortical sequence representation at the next position. We focused on position 3 in the sequence, as this is a critical decision point where one must choose amongst three possible states. Thus, we expected that hippocampal predictions about future states may be enhanced at this decision point (Johnson and Redish, 2007; Pfeiffer and Foster, 2013; Singer and Frank, 2009). If so, representations in the hippocampus at position 3 should share information with the patterns in cortex at position 4, where item representations and expectation effects are seen (Figure 4A).

**Figure 4.**
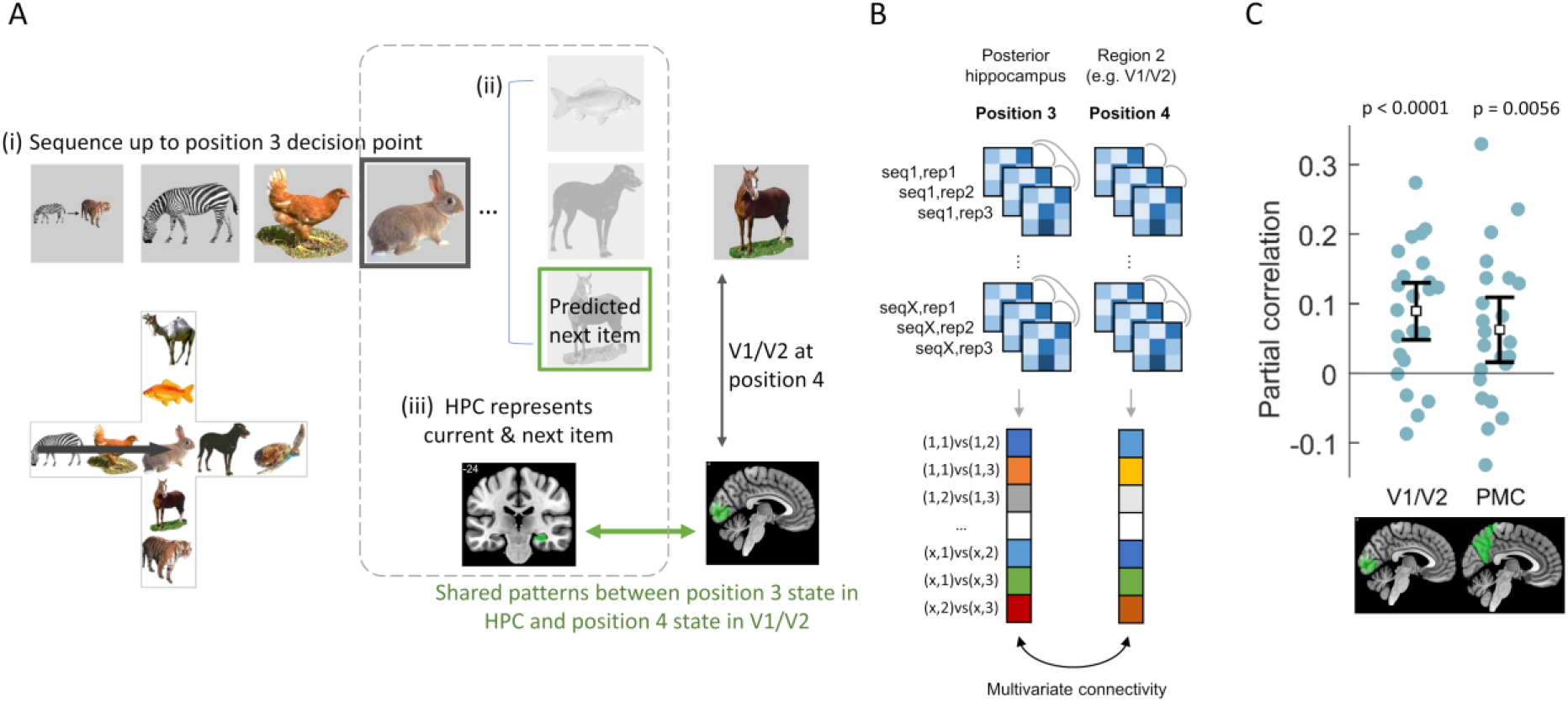
Cross region and position multivariate connectivity. A. (i) Participants navigate the sequence up to position 3, the central decision point of the cross-maze. (ii) From here it is possible to see 3 different animals, one of which is the correct next item for this sequence. (iii) If hippocampal representations are predictive of future states, then the hippocampal representation at position 3 will also contain some information about the correct future state (position 4). If so, representations in the hippocampus at position 3 should share information with the patterns in cortex at position 4, where item representations and expectation effects are seen. This is tested using multivariate connectivity. B) First, posterior hippocampal activity patterns are extracted for each rabbit item (position 3) for each repetition of each sequence. Pattern similarity is calculated between each repetition of the same-sequence pairs, resulting in a correlation vector, analogous to an unwrapped representational similarity matrix, but limited to same-sequence pairs. Following this, similarity is calculated between same sequence/item-pairs at position 4 for repetitions from the same sequence taken from a second region (e.g. V1/V2), producing a second similarity vector. Multivariate connectivity is calculated as the partial correlation between the two similarity vectors from the two regions/positions while controlling for the similarity of region 2 at position 3. C) Posterior hippocampus was significantly correlated with later similarity patterns in V1/V2 and PMC. Error bars show 95% CI around the mean.

To test this, we first calculated hippocampal PS between same sequence pairs at position 3, which is the central decision point in the sequence (where all items are rabbits; Figure 4B). We next calculated PS in cortical ROIs, V1/V2 and PMC, in all same sequence trial pairs at position 4. These ROIs were specifically chosen as they showed item expectation effects, suggesting they are a target for item reactivation. In order to provide control over auto-correlated effects between position 3 and 4 within a single region, we calculated the partial correlation between position 3 posterior hippocampal PS and position 4 PS from cortical regions while controlling for position 3 PS from the same cortical region. This analysis thus allows us to ask whether information in the hippocampus at position 3 is related to cortical information at the next point in the sequence. We find significant partial correlations between position 3 representations in posterior hippocampus and position 4 representations in V1/V2 (mean p_*xy*_ = 0.089, t(22) = 4.51, p < 0.0001) and PMC (mean p_*xy*_ = 0.063, t(22) = 2.77, p = 0.0056; Figure 4C). These results show that hippocampal responses reflect information about the upcoming states that later emerge in early visual and PMC, and is consistent with a predictive role of the hippocampus in supporting cortical reinstatement of expected future items.

## Discussion

In this study, we sought to uncover how the hippocampus, PM network and early visual cortex interact to support the prediction and reactivation of sensory details in cortex. Our results revealed that the human hippocampus and posterior cortical regions – the posterior medial cortex (precuneus/posterior cingulate cortex) and early visual regions, work together to represent, predict and reactivate expected sensory details of future events. After learning the locations of nine animals within a cross maze structure, participants moved through the maze during fMRI where they saw predictable sequences of animal images. Importantly, in our paradigm, one quarter of the sequences ended early, meaning that the next image in the sequence was expected but not shown, allowing us to investigate the nature of top-down expectations in the absence of any visual input. Pattern similarity (PS) analysis revealed complementary roles for the hippocampus and neocortical areas during navigation. Specifically, the posterior hippocampus carried information about items and sequence contexts, whereas visual cortical areas showed significant representations of the currently processed item, as well as predicted items that were not shown in a sequence. Furthermore, we found that the fidelity of hippocampal PS predicted subsequent item-specific representations in early visual cortex and the PMC. These findings show that hippocampal representations are used to generate expectations of future inputs via top-down modulation to the neocortex.

### Hippocampal memories guide the reactivation of upcoming sensory details

Using multivariate connectivity, we tested if the representational similarity structure seen in the hippocampus was related to representational similarity seen in V1/V2 and PMC, with the inference being that correlated representational states between regions indicates representations are shared between the regions (Pillet et al., 2020). Importantly, we calculated multivariate connectivity between the hippocampus at position 3 in the sequence (a rabbit image in all sequences) and V1/V2 and PMC at position 4, meaning that in addition to testing for shared representations across regions, we further tested the idea that information is shared between *past* hippocampal responses and *future* cortical responses. Using this approach, we observed a significant relationship between the posterior hippocampus at position 3 with both V1/V2 and PMC at position 4. These results argue that the hippocampus is a top-down source of predictive effects, which are then reactivated in cortex. Note that this effect cannot be driven by a concurrent strong representation of the current item in both regions, as our analysis controls for effects in the cortical regions at position 3. This means there is some element of representational similarity in the hippocampus at position 3 that can explain future representational similarity seen at position 4 in cortex, over and above that explained by the cortical regions at position 3 themselves. These results align with the view that hippocampal memories guide prediction of upcoming sensory events.

How might these predictive effects come about? One line of research to illustrate the link between the hippocampus and learning and prediction, is that of statistical learning. Studies of statistical learning argue that the hippocampus enables us to learn the structure of our environment, which can then be used to predict upcoming events to help guide behaviour (Kourtzi and Welchman, 2019; Schapiro et al., 2014; Sherman et al., 2020; Turk-Browne, 2019). Human fMRI data indeed points to the hippocampus for representing the temporal order of learned object sequences (Hsieh et al., 2014) and for predicting future states during navigation in learned environments (Brown et al., 2016).

Further, hippocampal place cells have been shown to reactivate prospective future locations along a navigational path, a phenomenon termed preplay (Johnson and Redish, 2007; Lisman and Redish, 2009). Such hippocampal predictions are also captured through differing computational accounts, that predict future states based on the current state and past learned experiences (Mattar and Daw, 2018; Stachenfeld et al., 2017). Regardless, together with our results, this points to a mechanism whereby the hippocampus is engaged in predicting upcoming events, through reinstating learned details.

Learning paradigms have further been employed to reveal the instantiation of visual predictions, where after learning sequences of visual gratings, the orientation of an expected grating can be decoded from early visual cortex (Luft et al., 2015). In conjunction with our data, these studies point to the hippocampus being a source of top-down modulation on early visual regions. Predictions about upcoming items could be reactivated in the hippocampus, through pattern completion (McClelland et al., 1995), with information about the expected sensory details then reactivated in cortex. In humans, evidence is emerging linking hippocampal pattern completion to visual predictions in sensory regions (Hindy et al., 2016; Kok and Turk-Browne, 2018). For example, Hindy et al., (2016) used high resolution fMRI after participants learned sequences of cue-response-outcome associations. Using multivariate classifiers trained on either the full sequence (cue-response-outcome) or the outcome alone, and applied to cue-response trials, they showed that hippocampal subfields CA1 and CA2/3DG contained information about the full sequence, while V1 and V2 contained information about an expected perceptual outcome. Further, Hindy et al., (2016) showed that sequence decoding in the hippocampus was related to outcome decoding in visual cortex, providing evidence that hippocampal responses were linked to the reactivation of predicted sensory details in visual cortex. These results parallel our hippocampal sequence effects and early visual cortex item expectation effects. However, much of the previous evidence focussed only on the hippocampus and visual cortex, while prediction was also part of the task. Here, participants navigated through learned environments which resulted in seeing sequences of visual objects in a task that does not overtly emphasise prediction. This allowed us to further establish the top-down nature of hippocampal representations with visual cortex, but we also extend past research by showing top-down effects in the contextually sensitive posterior medial cortex, and during a task that involved goal-directed navigation, rather than cue-outcome predictions. Our data further adds to a broad literature highlighting a top-down modulatory role of the human hippocampus on visual cortex, and beyond, during memory-guided behaviours such as retrieval (Barron et al., 2020; Bosch et al., 2014; Staresina and Wimber, 2019; Warren et al., 2019), navigation (Bridge et al., 2017; Sherrill et al., 2013; Watrous and Ekstrom, 2014) and attention (Aly and Turk-Browne, 2016a, 2016b; Günseli and Aly, 2020).

### Predicted sensory details are reactivated in early visual and PM cortex

A critical question we addressed was which regions represented expected items in the absence of any bottom-up input, finding that both V1/V2 and the PMC showed item expectation effects. These findings were possible due to the catch trials included in the paradigm, where, after position 3 in the sequence, the final two items of the sequence were omitted and replaced by an extended blank screen. This meant that we could evaluate representations of what was expected, but did not appear. Such expectation effects require the retrieval of information based on past learned experiences with a given sequence context, and the reactivation of the expected sensory patterns.

Previous studies have reported the reactivation of expected sensory details in primary visual regions for abstract stimuli (Alink et al., 2010; Eagleman and Dragoi, 2012; Kok et al., 2014, 2012), with our results showing this effect generalises to more complex meaningful items. Going beyond these studies, our results revealed item expectation effects outside of primary visual cortex - in the PMC. The design of our study did not permit us to dissociate between visual and PMC representations, though the extant literature speaks to this issue. For example, recent fMRI research shows the PMC becomes more engaged when item predictions can be made from a preceding visual scene. Caplette et al., (2020) showed visual scenes followed by objects which were either predictable given general knowledge about a scene (e.g. a beachball is consistent with a beach scene) or the object was not predictable following a scrambled scene. They showed increased activity in the precuneus for objects that could be predicted, suggesting the precuneus is integrating information about the context specific predictions and the object. In our paradigm, like the early visual cortex, the PMC showed expectation effects when an item was expected, but never shown. Such expectation effects in the PMC have also been observed during speech following a predictive context (Scharinger et al., 2016). Such findings suggest that the PMC might play a role in prediction that transcends sensory modalities, consistent with the idea that PMC representations may be relatively abstract or semantic in nature (Ranganath and Ritchey, 2012).

The item expectation effects we see are driven by the differentiation of activity patterns to different expected animals. Several lines of evidence have shown that when an item is highly predictable, there is both an increase in stimulus decoding, and decrease in activity magnitude (Alink et al., 2010; Kok et al., 2012). The stimulus-specific nature of our expectation effects, observed without the occluding impact of perceiving a stimulus, are consistent with models where predictions result in a sharpening of neural representations, resulting in reduced BOLD signals, as a consequence of top-down constraints (de Lange et al., 2018). In our study, the reactivated patterns were specific enough to distinguish between different expected animal images. Yet what is less clear is the level of detail and nature of information that was reactivated. As we hypothesise above, it is likely that low-level visual details are reactivated in early visual areas, and higher-level visual, semantic and contextual signals in the PMC. In real life situations, our expectations are less constrained than in the current paradigm, and so predictions are unlikely to be related to a specific perceptual stimulus. Rather, real-world predictions must be more general than the stimulus-templates hypothesised to be reactivated in primary visual regions. As such, important future work should investigate how the specificity of mnemonic expectations modulates the strength and mechanism of hippocampal guided predictions.

It is long established that our past experiences impact our current perceptions. The current study provides novel insights into the mechanisms of how top-down expectations can influence the visual processing of objects. The current results advance our understanding of how the hippocampus and posterior cortical regions work together to support perceptual expectations and predictions of future states based on learned contexts. Further, our results help to bridge between research on the cortical manifestation of expectations, and research on predictive and contextual representations in the hippocampus.

## Methods

### Participants

Thirty healthy individuals participated in the study. All participants had normal or corrected-to-normal vision and were right handed. Data from one participant was excluded due to technical complications with the fMRI scanner, one subject was excluded due to incomplete behavioural data, two subjects were excluded due to poor behavioral performance in the scanner, and one subject was removed from the scanner before the experiment was completed. Prior to data analysis, to ensure data quality, we conducted a univariate analysis to look at motor and visual activation during the task compared to an implicit baseline. Two subjects showed little to no activation in these regions and were excluded from further analysis. This resulted in twenty-three participants reported here (11 male, 12 female, all right handed). Written informed consent was obtained from each participant prior to the experiment, and the study was approved by The Institutional Review Board at the University of California, Davis.

### Stimuli

The stimuli consisted of nine common animal images. Each animal was represented by a colour photograph, presented in isolation on a grey background. The nine animals were positioned into a cross maze with two animals per arm and one in the central location, creating the first zoo called ‘San Francisco Zoo’. A second zoo was created, ‘San Diego Zoo’, by mirror-reversing and rotating 90 degrees anti-clockwise one zoo map to create the other. Therefore, both zoo contexts contained the same animals with the same transitional structure but a different global layout.

### Procedure

#### Training

Participants initially underwent a training session in order to learn the animals and their locations within the two zoos. To achieve this, participants completed map construction, exploration and navigation trials for one zoo, with the process then repeated for the second (zoo order counter-balanced across participants). During map construction, participants were shown the zoo layout and asked to arrange the animal images shown on screen into the correct positions. During exploration, participants were shown the animal at the centre of the maze and were free to move up/down/left/right to see how moving in different directions resulting in seeing a different animal. After making 9 moves, the exploration reset to the centre animal to begin again. Exploration continued until each animal was seen at least four times. Next, participants navigated between positions in the maze. They were shown a cue image indicating a start and a goal animal, followed by seeing the start animal. The participant had to select the correct moves to reach the goal animal. Start and goal animals were always located at the end points of an arm, and participants had a maximum of four moves. If participants did not reach criterion during navigation, they repeated map construction and navigation. After completing training with one zoo, the procedure was repeated with the second zoo. After successfully completing training with both zoos, participants completed a final navigation task including trials from both zoos and presented with the same structure and timings of the fMRI navigation task (see below). All training and navigation tasks were presented using Psychtoolbox and Matlab.

#### fMRI

During fMRI scanning, participants performed the navigation task for both zoos in a blocked fashion across six scanning runs. In each run, participants were told which zoo they were in and completed eight navigation trials for one zoo before switching to the other zoo. A navigation trial consisted of seeing a cue screen showing the start and goal animal for 3 seconds, followed by a blank screen for 3 seconds. The start animal was then displayed for 2 seconds followed by a 3 second blank screen. After a response, the relevant animal was shown for 2 seconds followed by a 3 second blank screen, with the process continuing for a maximum of four moves. After four responses were made, the participant was shown a feedback screen. The participant was required to make their response within 2 seconds of the animal appearing, otherwise the move was judged as incorrect and they were shown a text screen indicating ‘wrong move’ for 2 seconds followed by a 3 second blank screen, and the animal was shown again.

Participants competed navigation trials for the twelve possible start and goal animal combinations in each zoo, with each navigation trial being repeated 3 times resulting in 72 full navigation trials. An additional 24 catch navigation trials were included where each navigation trial was terminated early, after the third animal (the central animal if correct responses were made), and instead of seeing the fourth animal, participants saw an additional blank screen lasting six seconds before a new trial began. The order of zoos was counterbalanced both across runs and between participants.

#### Scanning acquisition

MRI data were acquired on a 3T Siemens Skyra MRI using a 32-channel head coil. Anatomical images were collected using a T1-weighted magnetization prepared rapid acquisition gradient echo (MPRAGE) pulse sequence image (TR = 1800ms; TE = 29.6 ms; flip-angle = 7 degrees; 1 mm^3^ isotropic voxels; 208 axial slices, TR=2100ms, TE=2.98ms, FOV = 256mm). Functional images were collected with a multi-band gradient echo planar imaging sequence (TR = 1222 ms; TE = 24 ms; flip angle = 67 degrees; matrix=64×64, FOV=192mm; multi-band factor = 2; 3 mm^3^ isotropic spatial resolution).

#### Preprocessing

Preprocessing used SPM12 (https://www.fil.ion.ucl.ac.uk/spm/). Functional images underwent slice time correction, spatial realignment and smoothing using a 4mm FWHM Gaussian kernal. To detect fast motion events, the ART repair toolbox (Mazaika et al., 2009) was used. These spike events were used as nuisance variables within the GLMs. Single item beta images were obtained by running a separate GLM for each object (LSS model; Mumford et al., 2012). For each GLM, the item of interest was entered as a single regressor with 1 event, with an additional regressor for all other events. All events were modelled as a 2 second boxcar and convolved with a canonical HRF. Additional regressors were included for each spike event identified, 12 motion regressors (6 for realignment and 6 for the derivatives of each of the realignment parameters), and a drift term using a 128s cutoff. This resulted in five images per full navigation trial (e.g. zebra, chicken, rabbit, horse, tiger) and 4 images for each catch navigation trial (e.g. zebra, chicken, rabbit, omitted item).

### Pattern Similarity Analysis

#### ROI PS

Our initial analyses focussed on anatomical regions of interest (ROIs) in early visual cortex and the hippocampus. A V1/V2 region was created from the functional atlas of visual cortex developed by Rosenke et al. (2021), where V1 and V2 were combined into a single region, and inverse normalised to native space. Probabalistic maps of the hippocampal head, body and tail were obtained from the multistudy group template (Yushkevich et al., 2015). These maps were warped to MNI space using DARTEL and thresholded at 0.5. The resulting maps were then reverse normalized to each participants native space using Advanced Normalization Tools. The anterior hippocampus was defined as the hippocampal head, and the posterior hippocampus as the combined body and tail sections. This division closely follows recommended anterior–posterior divisions (Poppenk et al., 2013). In addition, ROIs of the perirhinal cortex (PRC) and parahippocampal cortex (PHC) were obtained from the multistudy group template (Yushkevich et al., 2015), and the temporal pole and PMC (combined precuneus and posterior cingulate cortex) were generated using FreeSurfer (version 6) and warped to each participants native space.

For item effects, pattern similarity was calculated using all grey matter voxels within each ROI and tested using a one-sampled t-test against zero. Item expectation effects were also tested in the ROIs showing significant item effects, and were based on pattern similarity between the expected-but-omitted beta image and the beta image of trials where the same item was seen. Specifically, pattern similarity was calculated between each omitted item and the position 4 trials where the same item was seen. Pattern similarity was then averaged across pairs where the omitted item was due to be the same as the seen item. This was contrasted with pattern similarity where the omitted item was different to the seen item. To test for effects of sequence, PS was calculated between pairs of trials that shared the same item and the same sequence context (ignoring the zoo context) and contrasted with PS for the same-items from a different sequence. All pattern similarity values were calculated between pairs of items using Pearson’s correlation, excluding items where an incorrect response was made and excluding pairs of trials that occurred in the same scanning run (Mumford et al., 2012).

#### Between region PS

We also tested the degree to which PS in one region related to PS in another region at a different position in the trial. Here, we tested whether PS in the hippocampus at position 3 in the sequence was related to PS at position 4 in cortical regions that showed item expectation effects. To do this, we used a partial correlation analysis whereby hippocampal PS from position 3 was correlated with position 4 PS from a second region, while controlling for PS from the second region at position 3. The analysis tells us if past information in the hippocampus can explain future information in cortex, over and above that explained by past information in that same cortical region. Our analysis focused on PS between trials from the same sequence, and same zoo (excluding across sequence/zoo PS).

## Acknowledgements

This research was funded by a Royal Society and Wellcome Trust Sir Henry Dale Fellowship to AC (211200/Z/18/Z) and U.S. Office of Naval Research Grants N00014-15-1-0033 and N00014-17-1-2961 to CR. Any opinions, findings, and conclusions or recommendations expressed in this material are those of the authors and do not necessarily reflect the views of the U.S. Department of Defense. For the purpose of open access, the author has applied a CC BY public copyright licence to any Author Accepted Manuscript version arising from this submission. The authors report no conflict of interest.

## References

Alink, A., Schwiedrzik, C.M., Kohler, A., Singer, W., Muckli, L., 2010. Stimulus Predictability Reduces Responses in Primary Visual Cortex. J. Neurosci. 30, 2960–2966. https://doi.org/10.1523/JNEUROSCI.3730-10.2010

Aly, M., Turk-Browne, N.B., 2016a. Attention promotes episodic encoding by stabilizing hippocampal representations. PNAS 113, E420–E429. https://doi.org/10.1073/pnas.1518931113

Aly, M., Turk-Browne, N.B., 2016b. Attention Stabilizes Representations in the Human Hippocampus. Cereb Cortex 26, 783–796. https://doi.org/10.1093/cercor/bhv041

Anzellotti, S., Coutanche, M.N., 2018. Beyond Functional Connectivity: Investigating Networks of Multivariate Representations. Trends in Cognitive Sciences 22, 258–269. https://doi.org/10.1016/j.tics.2017.12.002

Bar, M., 2009. The proactive brain: memory for predictions. Philosophical Transactions of the Royal Society B: Biological Sciences 364, 1235–1243. https://doi.org/10.1098/rstb.2008.0310

Bar, M., 2004. Visual objects in context. Nat Rev Neurosci 5, 617–29. https://doi.org/10.1038/nrn1476

Bar, M., Aminoff, E., 2003. Cortical analysis of visual context. Neuron 38, 347–58.

Barnett, A.J., Man, V., McAndrews, M.P., 2019. Parcellation of the Hippocampus Using Resting Functional Connectivity in Temporal Lobe Epilepsy. Front. Neurol. 0. https://doi.org/10.3389/fneur.2019.00920

Barnett, A.J., Reilly, W., Dimsdale-Zucker, H.R., Mizrak, E., Reagh, Z., Ranganath, C., 2021. Intrinsic connectivity reveals functionally distinct cortico-hippocampal networks in the human brain. PLOS Biology 19, e3001275. https://doi.org/10.1371/journal.pbio.3001275

Barron, H.C., Auksztulewicz, R., Friston, K., 2020. Prediction and memory: A predictive coding account. Progress in Neurobiology 192, 101821. https://doi.org/10.1016/j.pneurobio.2020.101821

Bosch, S.E., Jehee, J.F.M., Fernández, G., Doeller, C.F., 2014. Reinstatement of associative memories in early visual cortex is signaled by the hippocampus. J Neurosci 34, 7493–7500. https://doi.org/10.1523/JNEUROSCI.0805-14.2014

Bridge, D.J., Cohen, N.J., Voss, J.L., 2017. Distinct Hippocampal versus Frontoparietal Network Contributions to Retrieval and Memory-guided Exploration. Journal of Cognitive Neuroscience 29, 1324–1338. https://doi.org/10.1162/jocn_a_01143

Brown, T.I., Carr, V.A., LaRocque, K.F., Favila, S.E., Gordon, A.M., Bowles, B., Bailenson, J.N., Wagner, A.D., 2016. Prospective representation of navigational goals in the human hippocampus. Science 352, 1323–1326. https://doi.org/10.1126/science.aaf0784

Caplette, L., Gosselin, F., Mermillod, M., Wicker, B., 2020. Real-world expectations and their affective value modulate object processing. NeuroImage 213, 116736. https://doi.org/10.1016/j.neuroimage.2020.116736

Davenport, J.L., Potter, M.C., 2004. Scene consistency in object and background perception. Psychol Sci 15, 559–564. https://doi.org/10.1111/j.0956-7976.2004.00719.x

de Lange, F.P., Heilbron, M., Kok, P., 2018. How Do Expectations Shape Perception? Trends Cogn Sci 22, 764–779. https://doi.org/10.1016/j.tics.2018.06.002

Eagleman, S.L., Dragoi, V., 2012. Image sequence reactivation in awake V4 networks. PNAS 109, 19450–19455. https://doi.org/10.1073/pnas.1212059109

Eichenbaum, H., Fortin, N.J., 2009. The neurobiology of memory based predictions. Philos Trans R Soc Lond B Biol Sci 364, 1183–1191. https://doi.org/10.1098/rstb.2008.0306

Ezzyat, Y., Davachi, L., 2014. Similarity Breeds Proximity: Pattern Similarity within and across Contexts Is Related to Later Mnemonic Judgments of Temporal Proximity. Neuron 81, 1179–1189. https://doi.org/10.1016/j.neuron.2014.01.042

Günseli, E., Aly, M., 2020. Preparation for upcoming attentional states in the hippocampus and medial prefrontal cortex. eLife 9, e53191. https://doi.org/10.7554/eLife.53191

Hindy, N.C., Ng, F.Y., Turk-Browne, N.B., 2016. Linking pattern completion in the hippocampus to predictive coding in visual cortex. Nat Neurosci 19, 665–667. https://doi.org/10.1038/nn.4284

Hsieh, L., Gruber, M.J., Jenkins, L.J., Ranganath, C., 2014. Hippocampal activity patterns carry information about objects in temporal context. Neuron 81, 1165–1178.

Johnson, A., Redish, A.D., 2007. Neural Ensembles in CA3 Transiently Encode Paths Forward of the Animal at a Decision Point. J. Neurosci. 27, 12176–12189. https://doi.org/10.1523/JNEUROSCI.3761-07.2007

Jonker, T.R., Dimsdale-Zucker, H., Ritchey, M., Clarke, A., Ranganath, C., 2018. Neural reactivation in parietal cortex enhances memory for episodically linked information. PNAS 115, 11084–11089. https://doi.org/10.1073/pnas.1800006115

Kok, P., Failing, M.F., de Lange, F.P., 2014. Prior expectations evoke stimulus templates in the primary visual cortex. J Cogn Neurosci 26, 1546–1554. https://doi.org/10.1162/jocn_a_00562

Kok, P., Jehee, J.F.M., de Lange, F.P., 2012. Less Is More: Expectation Sharpens Representations in the Primary Visual Cortex. Neuron 75, 265–270. https://doi.org/10.1016/j.neuron.2012.04.034

Kok, P., Rait, L.I., Turk-Browne, N.B., 2019. Content-based Dissociation of Hippocampal Involvement in Prediction. Journal of Cognitive Neuroscience 32, 527–545.https://doi.org/10.1162/jocn_a_01509

Kok, P., Turk-Browne, N.B., 2018. Associative Prediction of Visual Shape in the Hippocampus. J. Neurosci. 38, 6888–6899. https://doi.org/10.1523/JNEUROSCI.0163-18.2018

Kourtzi, Z., Welchman, A.E., 2019. Learning predictive structure without a teacher: decision strategies and brain routes. Curr Opin Neurobiol 58, 130–134. https://doi.org/10.1016/j.conb.2019.09.014

Kriegeskorte, N., Mur, M., Bandettini, P., 2008. Representational similarity analysis - connecting the branches of systems neuroscience. Front Syst Neurosci 2, 4. https://doi.org/10.3389/neuro.06.004.2008

Lee, H., Kuhl, B.A., 2016. Reconstructing Perceived and Retrieved Faces from Activity Patterns in Lateral Parietal Cortex. J. Neurosci. 36, 6069–6082. https://doi.org/10.1523/JNEUROSCI.4286-15.2016

Lisman, J., Redish, A.D., 2009. Prediction, sequences and the hippocampus. Philos Trans R Soc Lond B Biol Sci 364, 1193–1201. https://doi.org/10.1098/rstb.2008.0316

Livne, T., Bar, M., 2016. Cortical Integration of Contextual Information across Objects. J Cogn Neurosci 28, 948–958. https://doi.org/10.1162/jocn_a_00944

Long, N.M., Kuhl, B.A., 2021. Cortical Representations of Visual Stimuli Shift Locations with Changes in Memory States. Current Biology 31, 1119–1126.e5. https://doi.org/10.1016/j.cub.2021.01.004

Luft, C.D.B., Meeson, A., Welchman, A.E., Kourtzi, Z., 2015. Decoding the future from past experience: learning shapes predictions in early visual cortex. J Neurophysiol 113, 3159–3171. https://doi.org/10.1152/jn.00753.2014

Mattar, M.G., Daw, N.D., 2018. Prioritized memory access explains planning and hippocampal replay. Nat Neurosci 21, 1609–1617. https://doi.org/10.1038/s41593-018-0232-z

Mazaika, P., Hoeft, F., Glover, G.H., Reiss, A.L., 2009. Methods and Software for fMRI Analysis for Clinical Subjects. Presented at the Human Brain Mapping.

McClelland, J., McNaughton, B., O’Reilly, R., 1995. Why there are complementary learning systems in the hippocampus and neocortex: Insights from the successes and failures of connectionist models of learning and memory. Psychological Review 102, 419–457.

Mumford, J.A., Turner, B.O., Ashby, F.G., Poldrack, R.A., 2012. Deconvolving BOLD activation in event-related designs for multivoxel pattern classification analyses. Neuroimage 59, 2636–2643. https://doi.org/10.1016/j.neuroimage.2011.08.076

Munneke, J., Brentari, V., Peelen, M.V., 2013. The influence of scene context on object recognition is independent of attentional focus. Front Psychol 4. https://doi.org/10.3389/fpsyg.2013.00552

Oliva, A., Torralba, A., 2007. The role of context in object recognition. Trends Cogn. Sci. (Regul. Ed.) 11, 520–527. https://doi.org/10.1016/j.tics.2007.09.009

Palmer, tephen E., 1975. The effects of contextual scenes on the identification of objects. Memory & Cognition 3, 519–526. https://doi.org/10.3758/BF03197524

Pfeiffer, B.E., Foster, D.J., 2013. Hippocampal place-cell sequences depict future paths to remembered goals. Nature 497, 74–79. https://doi.org/10.1038/nature12112

Pillet, I., Op de Beeck, H., Lee Masson, H., 2020. A Comparison of Functional Networks Derived From Representational Similarity, Functional Connectivity, and Univariate Analyses. Front. Neurosci. 0. https://doi.org/10.3389/fnins.2019.01348

Poppenk, J., Evensmoen, H.R., Moscovitch, M., Nadel, L., 2013. Long-axis specialization of the human hippocampus. Trends Cogn Sci 17, 230–240. https://doi.org/10.1016/j.tics.2013.03.005

Ranganath, C., Ritchey, M., 2012. Two cortical systems for memory-guided behaviour. Nat Rev Neurosci 13, 713–26. https://doi.org/10.1038/nrn3338

Ritchey, M., Cooper, R.A., 2020. Deconstructing the Posterior Medial Episodic Network. Trends Cogn Sci 24, 451–465. https://doi.org/10.1016/j.tics.2020.03.006

Rosenke, M., van Hoof, R., van den Hurk, J., Grill-Spector, K., Goebel, R., 2021. A Probabilistic Functional Atlas of Human Occipito-Temporal Visual Cortex. Cerebral Cortex 31, 603–619. https://doi.org/10.1093/cercor/bhaa246

Schapiro, A.C., Gregory, E., Landau, B., McCloskey, M., Turk-Browne, N.B., 2014. The Necessity of the Medial Temporal Lobe for Statistical Learning. J Cogn Neurosci 26, 1736–1747. https://doi.org/10.1162/jocn_a_00578

Scharinger, M., Bendixen, A., Herrmann, B., Henry, M.J., Mildner, T., Obleser, J., 2016. Predictions interact with missing sensory evidence in semantic processing areas. Human Brain Mapping 37, 704–716. https://doi.org/10.1002/hbm.23060

Sherman, B.E., Graves, K.N., Turk-Browne, N.B., 2020. The prevalence and importance of statistical learning in human cognition and behavior. Curr Opin Behav Sci 32, 15–20. https://doi.org/10.1016/j.cobeha.2020.01.015

Sherrill, K.R., Erdem, U.M., Ross, R.S., Brown, T.I., Hasselmo, M.E., Stern, C.E., 2013. Hippocampus and Retrosplenial Cortex Combine Path Integration Signals for Successful Navigation. J. Neurosci. 33, 19304–19313. https://doi.org/10.1523/JNEUROSCI.1825-13.2013

Singer, A.C., Frank, L.M., 2009. Rewarded outcomes enhance reactivation of experience in the hippocampus. Neuron 64, 910–921. https://doi.org/10.1016/j.neuron.2009.11.016

Stachenfeld, K.L., Botvinick, M.M., Gershman, S.J., 2017. The hippocampus as a predictive map. Nature Neuroscience 20, 1643–1653. https://doi.org/10.1038/nn.4650

Staresina, B.P., Wimber, M., 2019. A Neural Chronometry of Memory Recall. Trends in Cognitive Sciences 23, 1071–1085. https://doi.org/10.1016/j.tics.2019.09.011

Summerfield, C., Egner, T., 2009. Expectation (and attention) in visual cognition. Trends Cogn. Sci. (Regul. Ed.) 13, 403–409. https://doi.org/10.1016/j.tics.2009.06.003

Trapp, S., Bar, M., 2015. Prediction, context, and competition in visual recognition. Ann. N.Y. Acad. Sci. 1339, 190–198. https://doi.org/10.1111/nyas.12680

Turk-Browne, N.B., 2019. The hippocampus as a visual area organized by space and time: A spatiotemporal similarity hypothesis. Vision Research 165, 123–130. https://doi.org/10.1016/j.visres.2019.10.007

Võ, M.L.-H., Boettcher, S.E., Draschkow, D., 2019. Reading scenes: how scene grammar guides attention and aids perception in real-world environments. Current Opinion in Psychology, Attention & Perception 29, 205–210. https://doi.org/10.1016/j.copsyc.2019.03.009

Warren, K.N., Hermiller, M.S., Nilakantan, A.S., Voss, J.L., 2019. Stimulating the hippocampal posterior-medial network enhances task-dependent connectivity and memory. Elife 8, e49458. https://doi.org/10.7554/eLife.49458

Watrous, A.J., Ekstrom, A.D., 2014. The Spectro-Contextual Encoding and Retrieval Theory of Episodic Memory. Front. Hum. Neurosci. 8. https://doi.org/10.3389/fnhum.2014.00075

Yushkevich, P.A., Pluta, J.B., Wang, H., Xie, L., Ding, S.-L., Gertje, E.C., Mancuso, L., Kliot, D., Das, S.R., Wolk, D.A., 2015. Automated volumetry and regional thickness analysis of hippocampal subfields and medial temporal cortical structures in mild cognitive impairment. Human Brain Mapping 36, 258–287. https://doi.org/10.1002/hbm.22627

