## Supplementary Information for "Contextual expectations shape cortical reinstatement of sensory representations"

### Supplementary materials

#### Supplementary methods

A searchlight multivariate pattern similarity analysis was performed to complement and confirm our ROI findings. At each voxel, activity values were extracted from grey matter voxels within a 7 mm radius. Pattern similarity was calculated between all pairs of items using Pearson's correlation, excluding items where an incorrect response was made and excluding pairs of trials that occurred in the same scanning run (Mumford et al., 2012). To calculate item effects, pattern similarity values were averaged across repetitions of the same item in the same navigation sequence and the same context, and contrasted with pattern similarity values between different items from the same navigation sequence and the same context. By keeping the navigation sequence and context constant, the effects of sequence and context are controlled. The item effect value was mapped back to the voxel at the centre of the searchlight, before repeating at each grey matter voxel.

The item effect map for each participant was normalised to the MNI template and spatially smoothed using a 6 mm FWHM Gaussian kernel. Item effect maps were entered into a random effects analysis (RFX) in SPM12 where each voxel was tested using a one-sampled t-test against zero. Voxelwise multiple comparisons correction was applied through a threshold of $p < 0.05$ FWE-corrected. Searchlight results are displayed using BrainNet viewer (Xia et al., 2013).

#### Supplementary results

Our main analysis relied on anatomically defined ROIs. To determine if there were any additional regions sensitive to item information, a searchlight analysis was performed contrasting PS for repetitions of the same item with PS between different items from the same sequence and zoo. Searchlight analysis identified four clusters showing greater PS for same items compared to different items, being bilateral clusters spanning from the occipital pole to the posterior fusiform gyrus and lateral occipitotemporal cortex, and bilateral clusters centred on the precentral gyrus (Figure S1A, Table S1). While the occipitotemporal cortex (OTC) cluster overlaps with V1/V2, it is more extensive along the ventral visual pathway reflecting the processing of visual objects beyond low-level details. Item effects were also present in the precentral gyrus (PG), however this may reflect the shared motor response required when seeing the same item in the same context. To examine the impact of a shared motor response, regional PS was further calculated for trials sharing the same button press and compared to PS for trials with different button presses – while discounting same item trials. This analysis revealed a highly significant effect of 'move' in PG (mean = 0.029, $t(22) = 5.54$, $p < 0.0001$). In addition, no such effects were seen in OTC, or any other ROI examined (all p's > 0.05), suggesting that item effects in other regions do not reflect the button press.

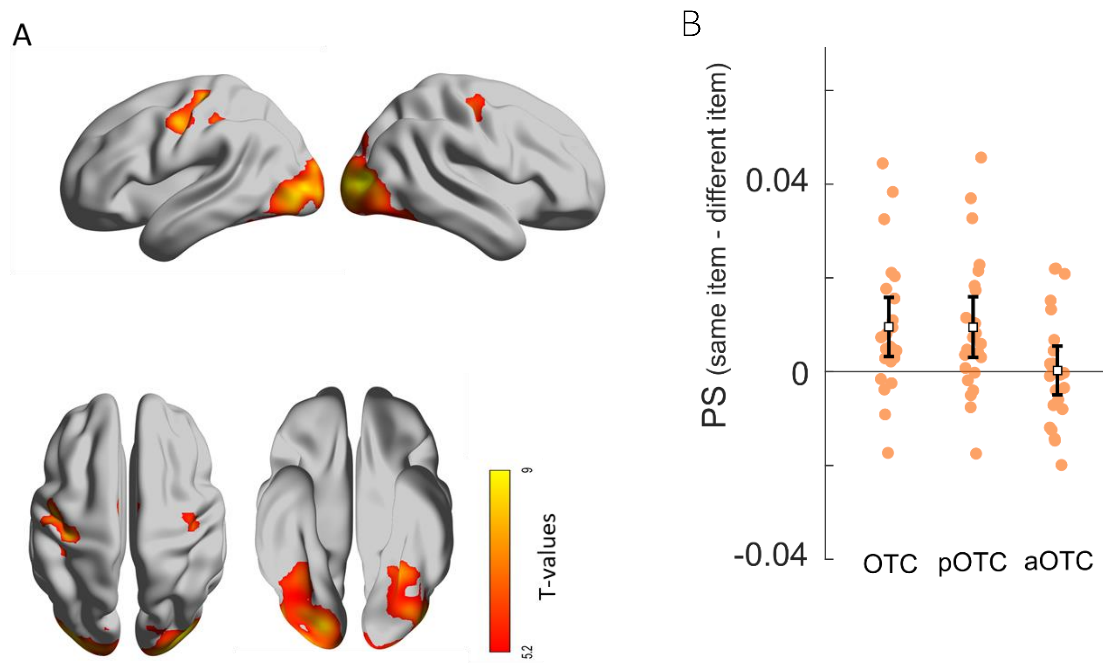


Figure S1. Item and expectation effects in visual cortex. A. Searchlight PS results showing higher PS for same items compared to different items within the same sequence and zoo context (voxel-wise FWE p < 0.05) resulting in two bilateral clusters in occipito-temporal cortex and the precentral gyrus. B. Item expectation effects. Error bars show 95% confidence intervals around the mean.

*Table S1. Searchlight results for item effects*

| Region | Cluster size | Location | T | p (FWE) | x | Y | Z |
|---|---|---|---|---|---|---|---|
| Occipitotemporal cortex | 1240 | R lateral occipital cortex | 11.14 | p < 0.001 | 33 | -82 | 5 |
| | | R occipital fusiform gyrus | 10.62 | p < 0.001 | 24 | -85 | 2 |
| | | R temporal occipital fusiform cortex | 7.84 | p = 0.001 | 36 | -49 | -28 |
| | 1094 | L lateral occipital cortex | 10.11 | p < 0.001 | -30 | -85 | -1 |
| | | L lingual gyrus | 8.08 | p = 0.001 | -14 | -55 | -16 |
| | | L temporal occipital fusiform cortex | 8.07 | p = 0.001 | -27 | -52 | -13 |
| Precentral gyrus | 712 | L precentral gyrus | 8.93 | p < 0.001 | -30 | -19 | 50 |
| | | L postcentral gyrus | 8.66 | p < 0.001 | -36 | -25 | 56 |
| | | L precentral gyrus | 8.28 | p < 0.001 | -45 | -10 | 44 |
| | 68 | R precentral gyrus | 6.79 | p = 0.007 | 36 | -16 | 53 |

*Locations from the Harvard-Oxford Atlas. Local maxima statistics shown after voxel-wise correction of p < 0.05 (FWE)*

We next tested whether the OTC showed item expectation effects based on similarity between the perceived trials and catch trials. Significant item expectation effects were seen in OTC (mean = 0.010, $t(22) = 3.19$, $p = 0.0024$; Figure S1B). Incorporating our earlier analysis of V1/V2, we have seen item expectation effects in both V1/V2 and a more extensive OTC region. This OTC region encompasses early visual areas as well as lateral occipital and posterior ventral temporal regions, and we next determined if expectation effects impacted only part, or all, of the OTC region. The OTC was split into a posterior section (occipital pole and lateral occipital cortex; pOTC) and anterior section (posterior fusiform; aOTC), after which significant item expectation effects were seen in the pOTC (mean = 0.010, $t(22) = 3.10$, $p = 0.003$) but not the aOTC (mean = 0.0002, $t(22) = 0.1$, $p = 0.46$), with significantly stronger effects in pOTC compared to aOTC ($t(22) = 2.97$, $p = 0.007$; Figure S1B). Therefore, our results point to visual expectation effects being predominantly concerned with lower-level visual regions in paradigms such as this where the exact stimulus can be predicted.

#### References:


Mumford, J. A., Turner, B. O., Ashby, F. G., & Poldrack, R. A. (2012). Deconvolving BOLD activation in event-related designs for multivoxel pattern classification analyses. *Neuroimage*, *59*(3), 2636–2643. https://doi.org/10.1016/j.neuroimage.2011.08.076

Xia, M., Wang, J., & He, Y. (2013). BrainNet Viewer: A Network Visualization Tool for Human Brain Connectomics. *PLOS ONE*, *8*(7), e68910. https://doi.org/10.1371/journal.pone.0068910
